## Supplementary figures and images for "Structure of the Calvin-Benson-Bassham sedoheptulose-1,7-bisphosphatase from the model microalga *Chlamydomonas reinhardtii*"

### supplementary figure 1

#### Slide 1
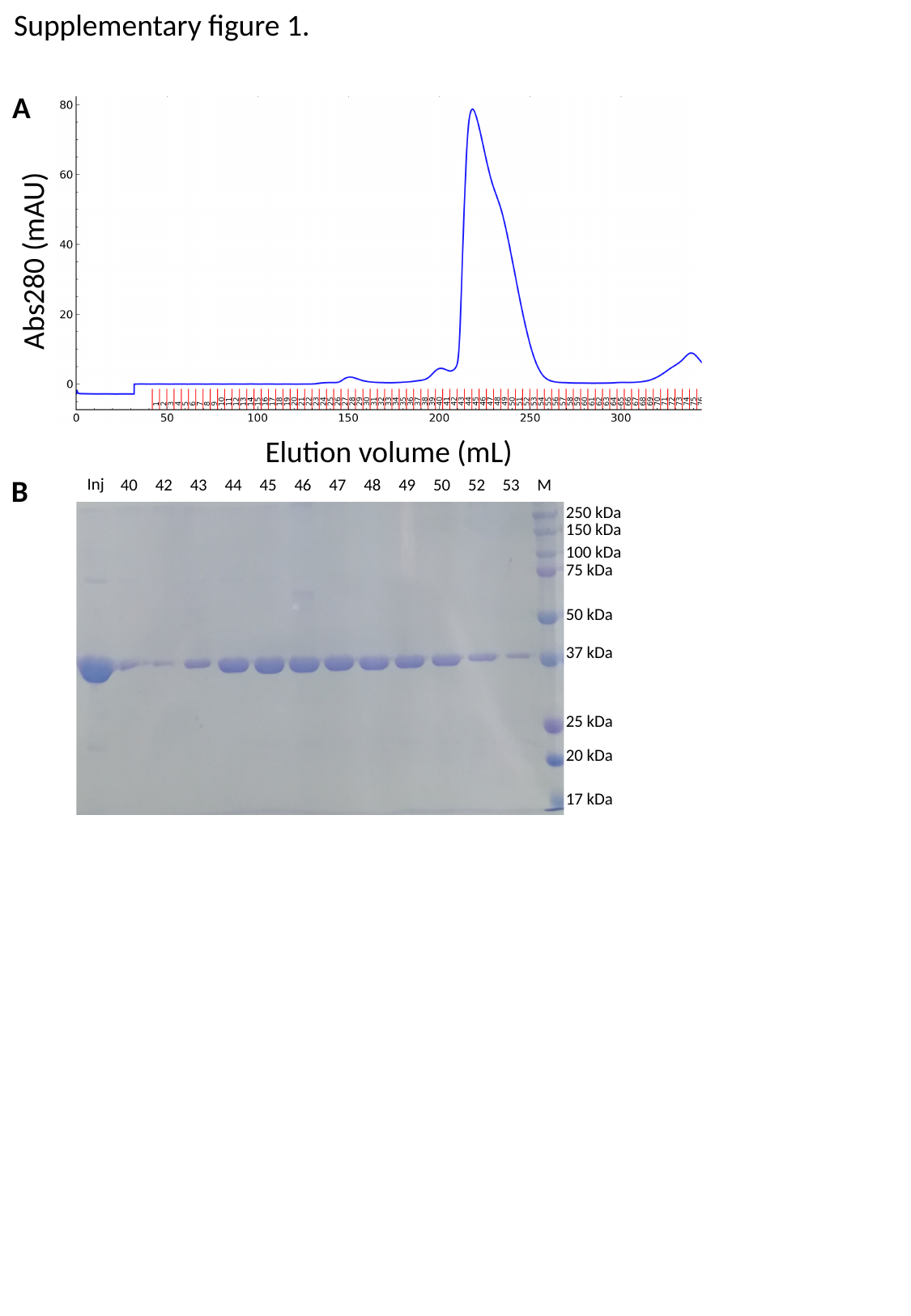

Supplementary figure 1.
A
Abs280 (mAU)
Elution volume (mL)
B
Inj
M
40
42
43
44
45
46
47
48
49
50
52
53
250 kDa
150 kDa
100 kDa
75 kDa
50 kDa
37 kDa
25 kDa
20 kDa
17 kDa

### supplementary figure 2

#### Slide 1
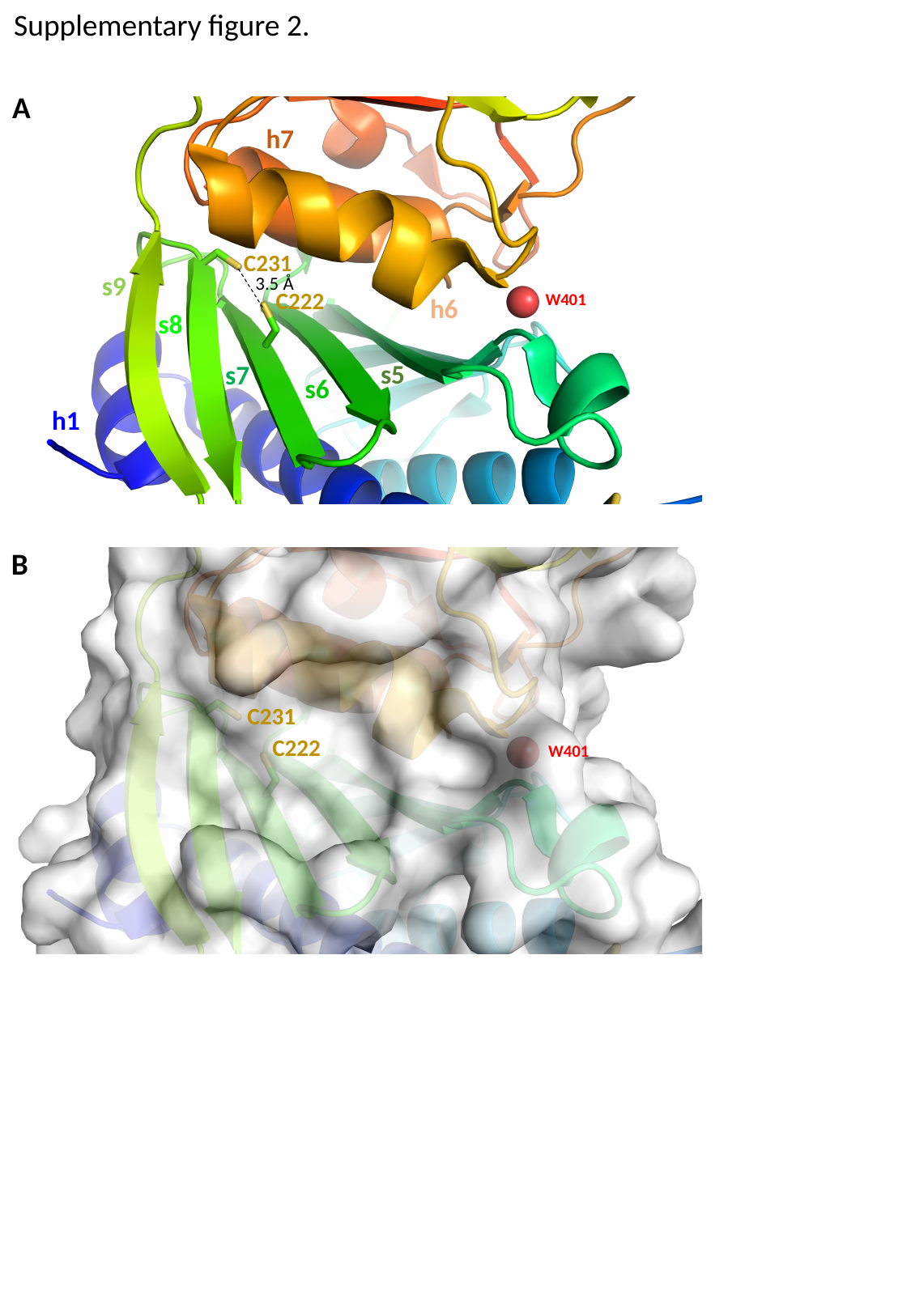

Supplementary figure 2.
A
h7
C231
s9
3.5 Å
C222
W401
h6
s8
s5
s7
s6
h1
B
C231
C222
W401

### supplementary figure 3

#### Slide 1
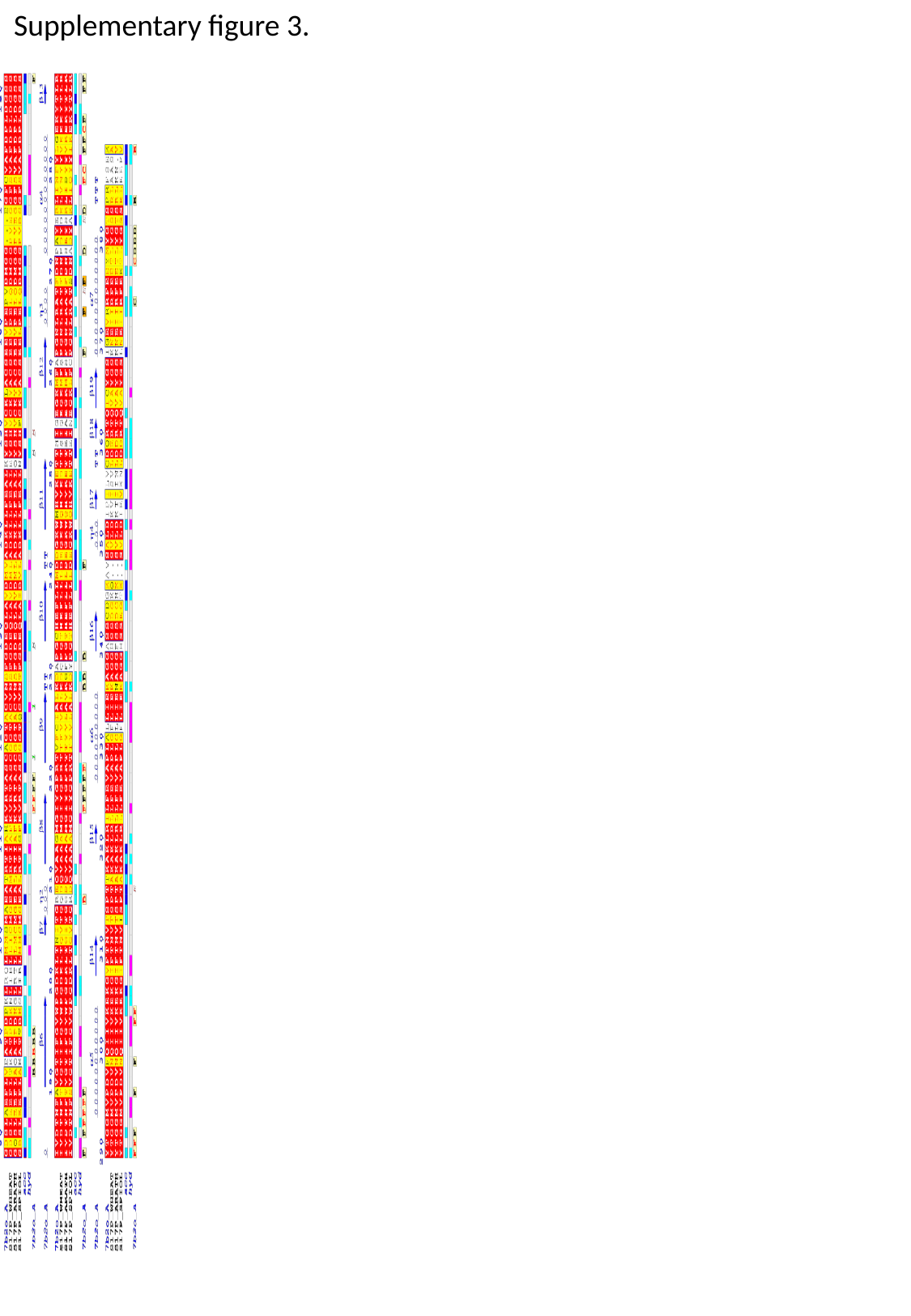

Supplementary figure 3.

### supplementary figure 4

#### Slide 1
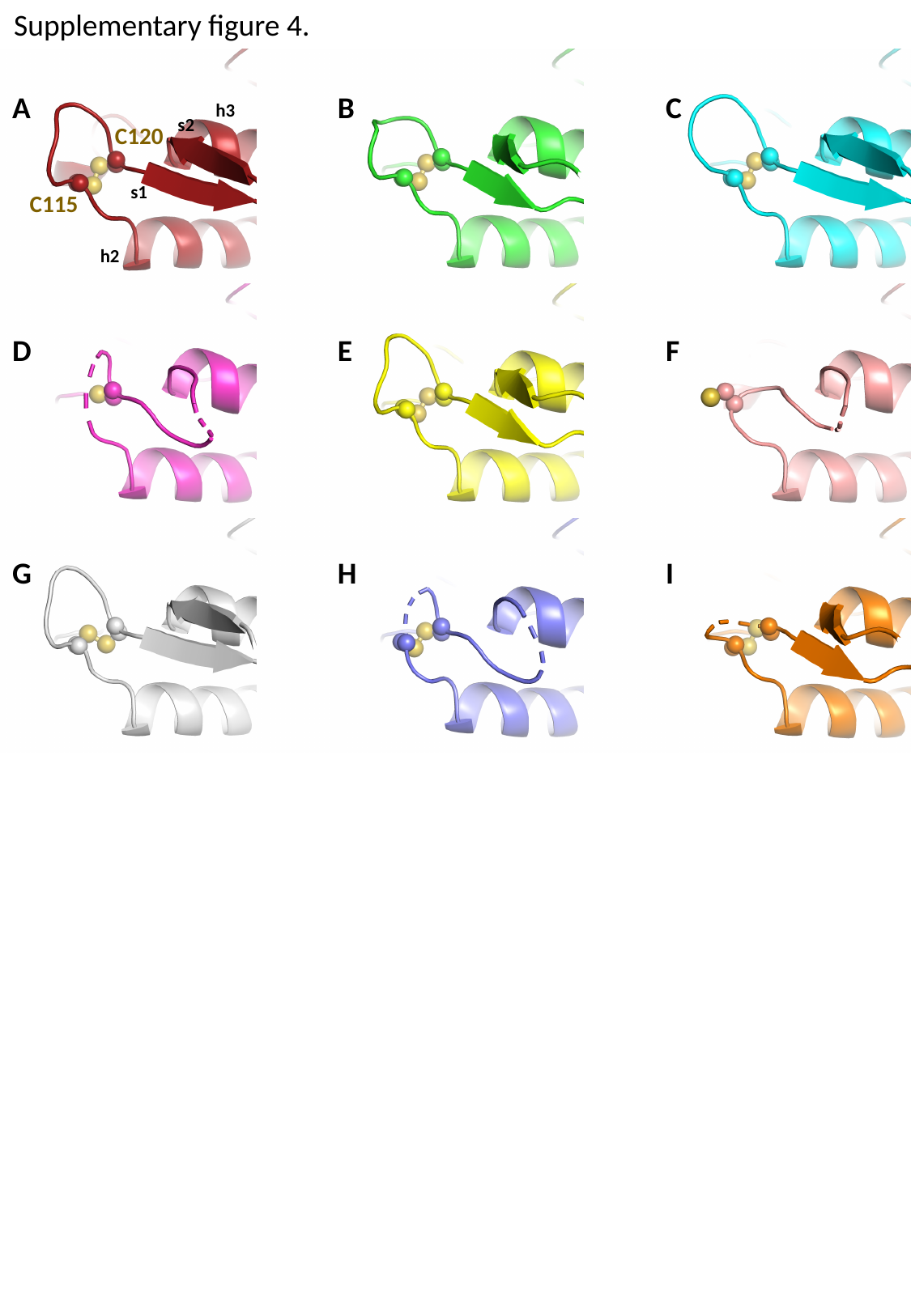

Supplementary figure 4.
A
B
C
h3
s2
C120
s1
C115
h2
D
E
F
G
H
I

### supplementary figure 6

#### Slide 1
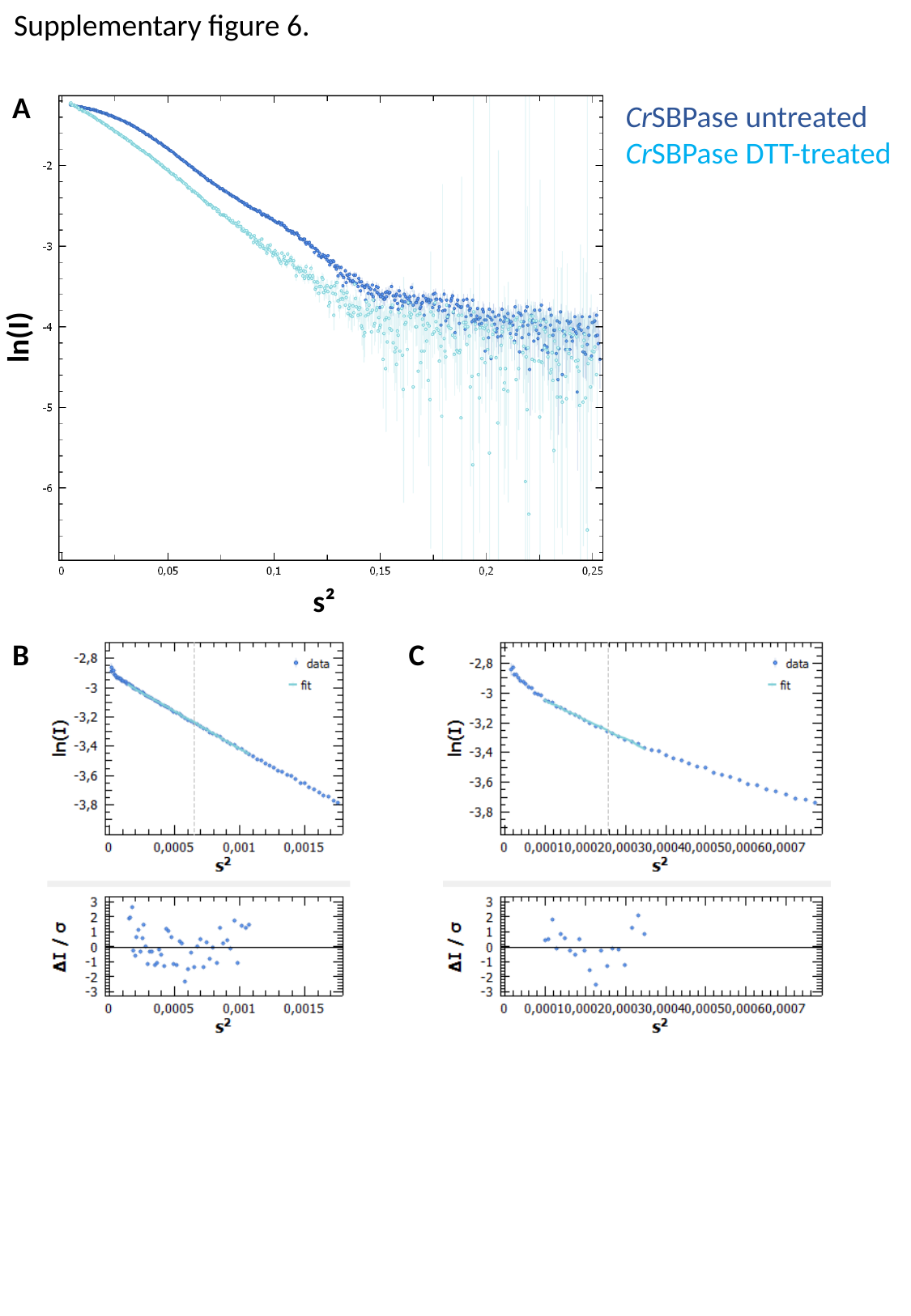

Supplementary figure 6.
A
CrSBPase untreated
CrSBPase DTT-treated
ln(I)
s²
B
C

### supplementary figure 7

#### Slide 1
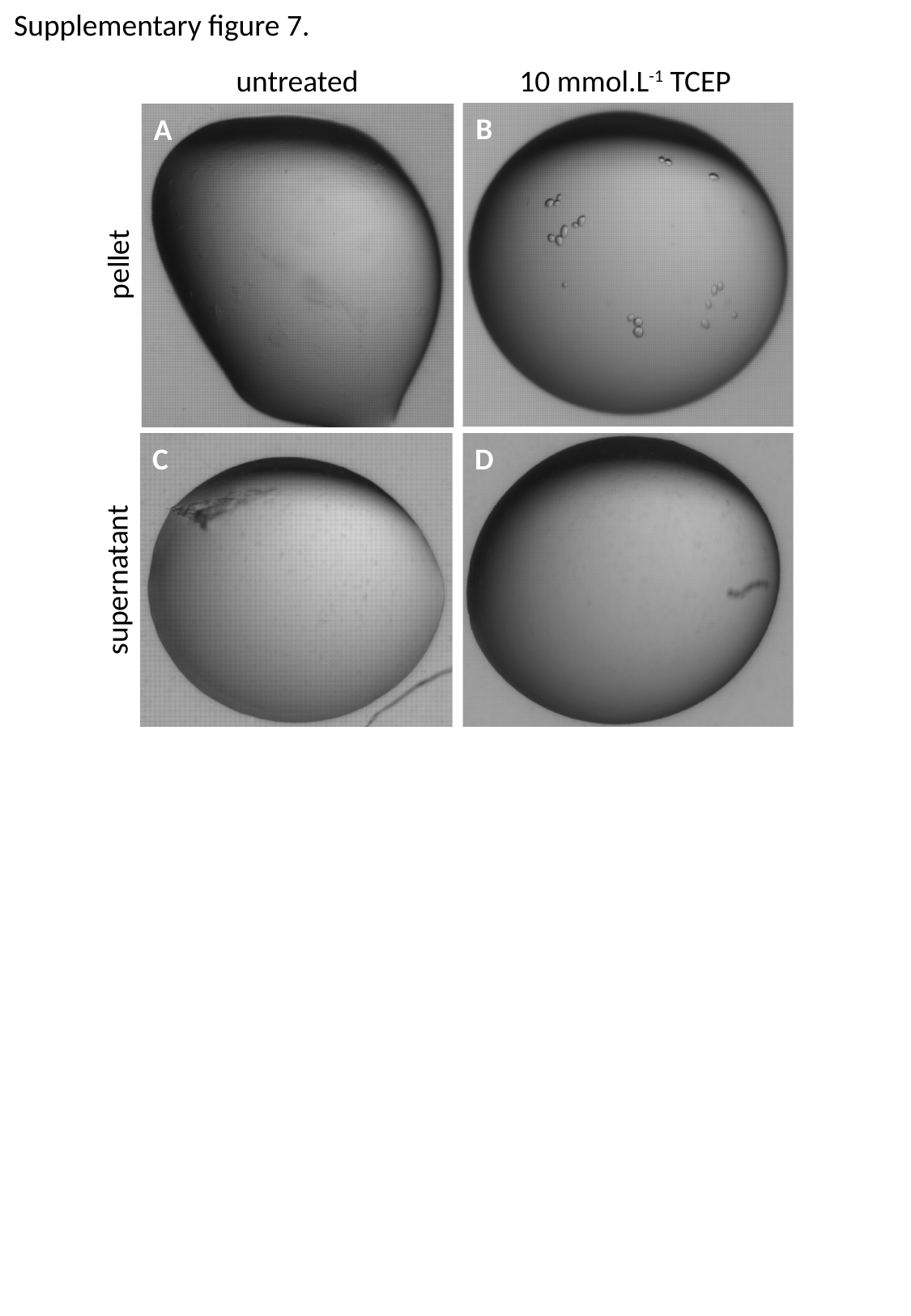

Supplementary figure 7.
untreated
10 mmol.L^-1^ TCEP
B
A
pellet
C
D
supernatant

### supplementary figure 8

#### Slide 1
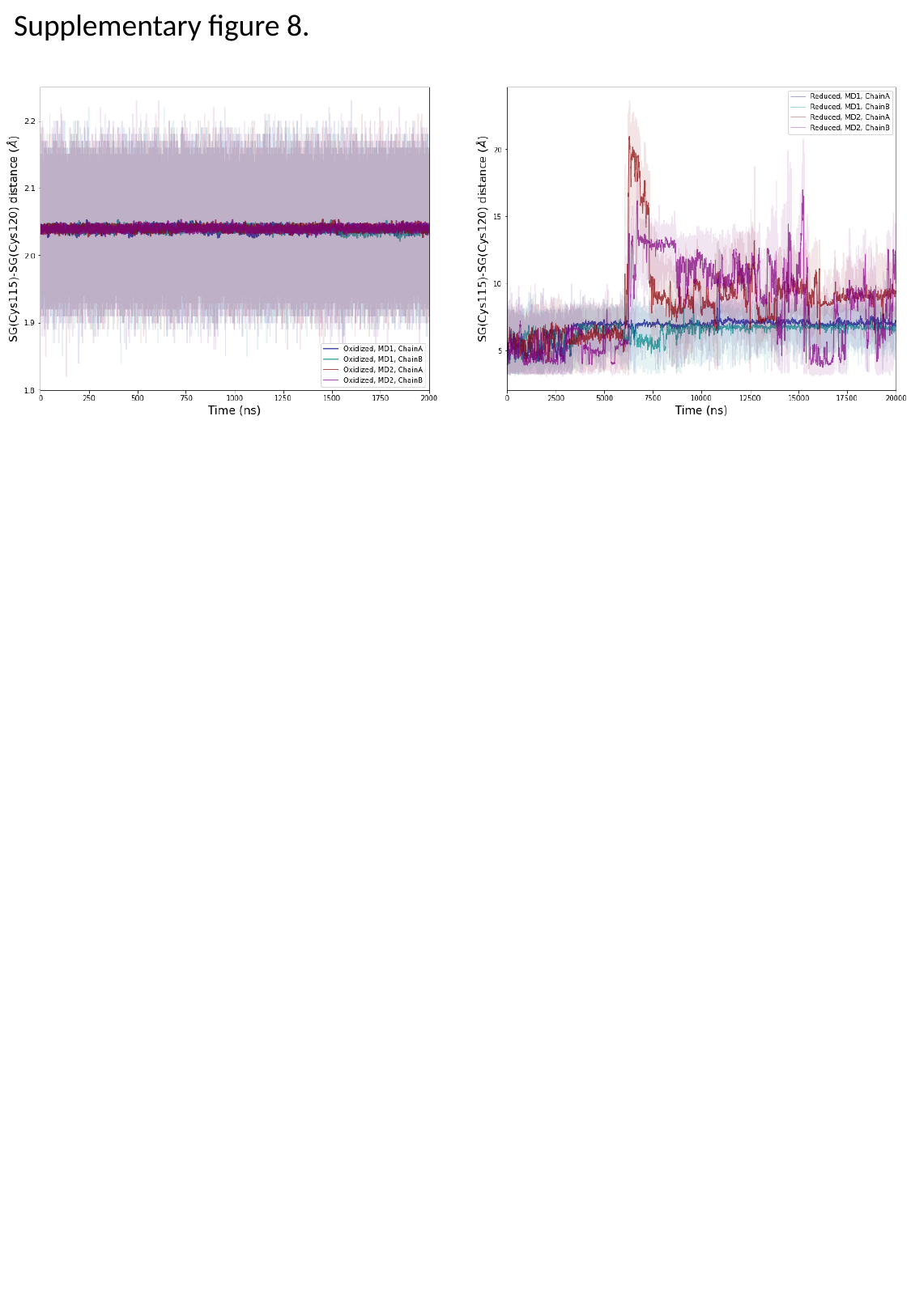

Supplementary figure 8.

### supplementary figure 9

#### Slide 1
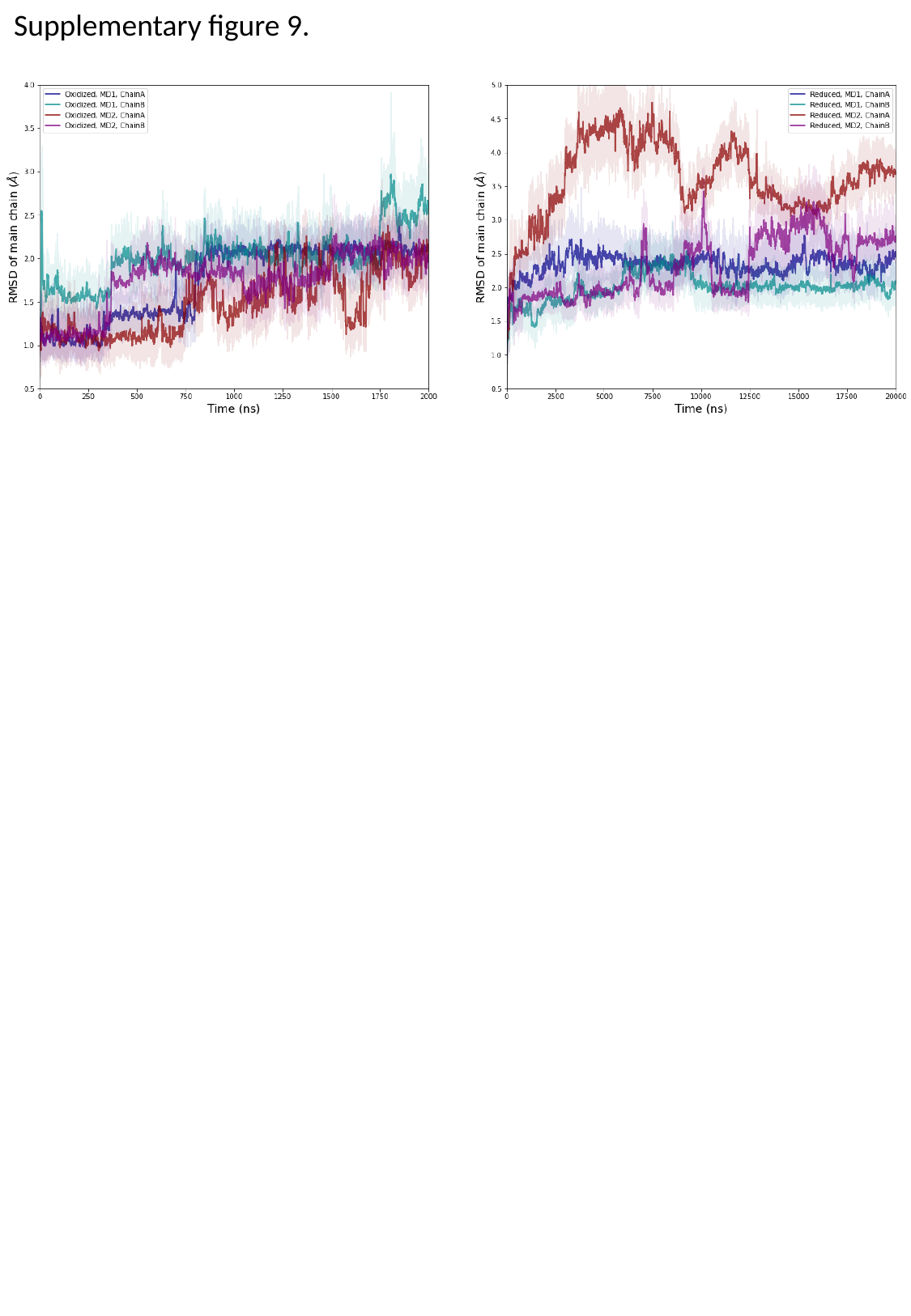

Supplementary figure 9.

### supplementary figure 10

#### Slide 1
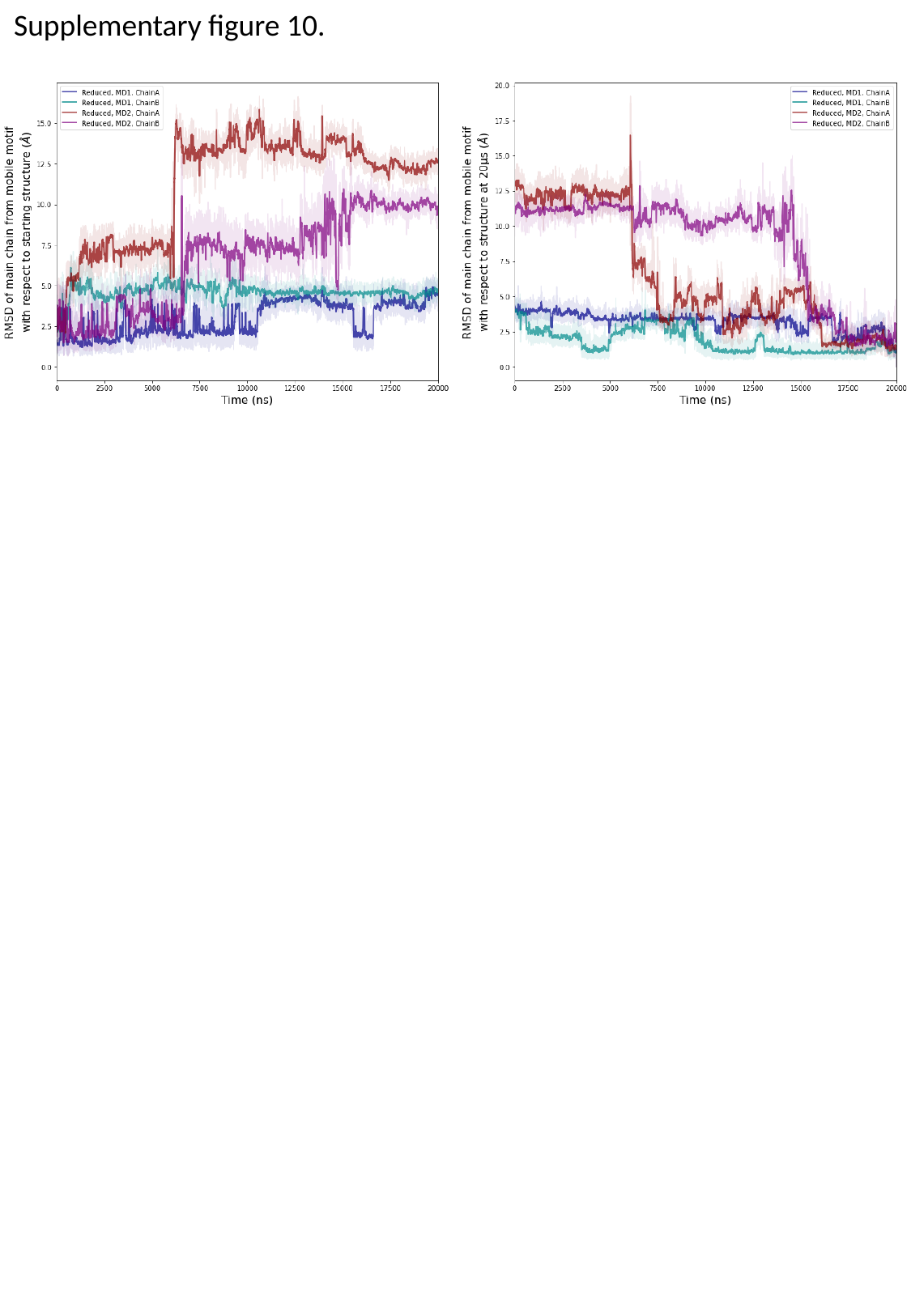

Supplementary figure 10.

### supplementary figure 11

#### Slide 1
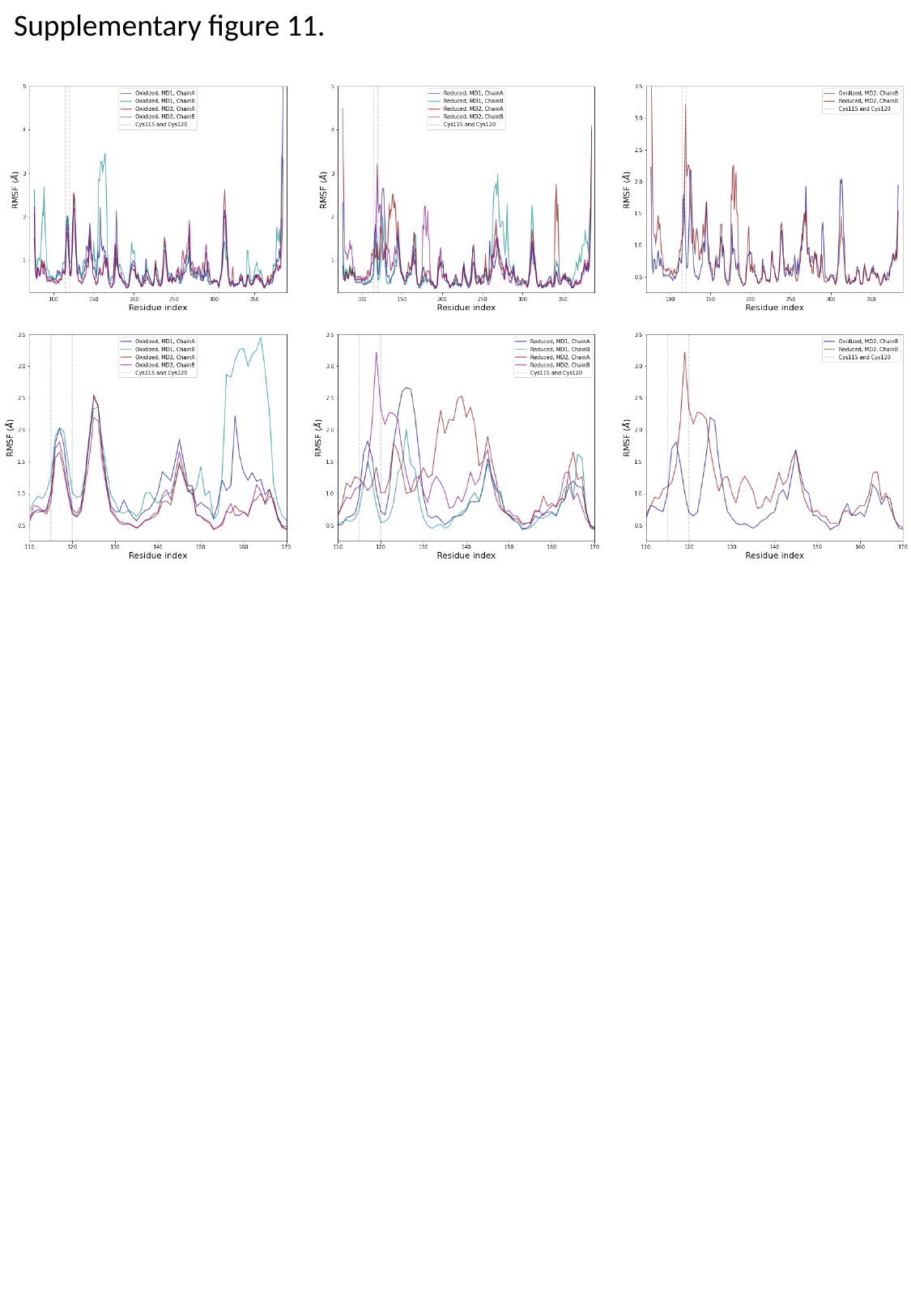

Supplementary figure 11.

### supplementary figure 12

#### Slide 1
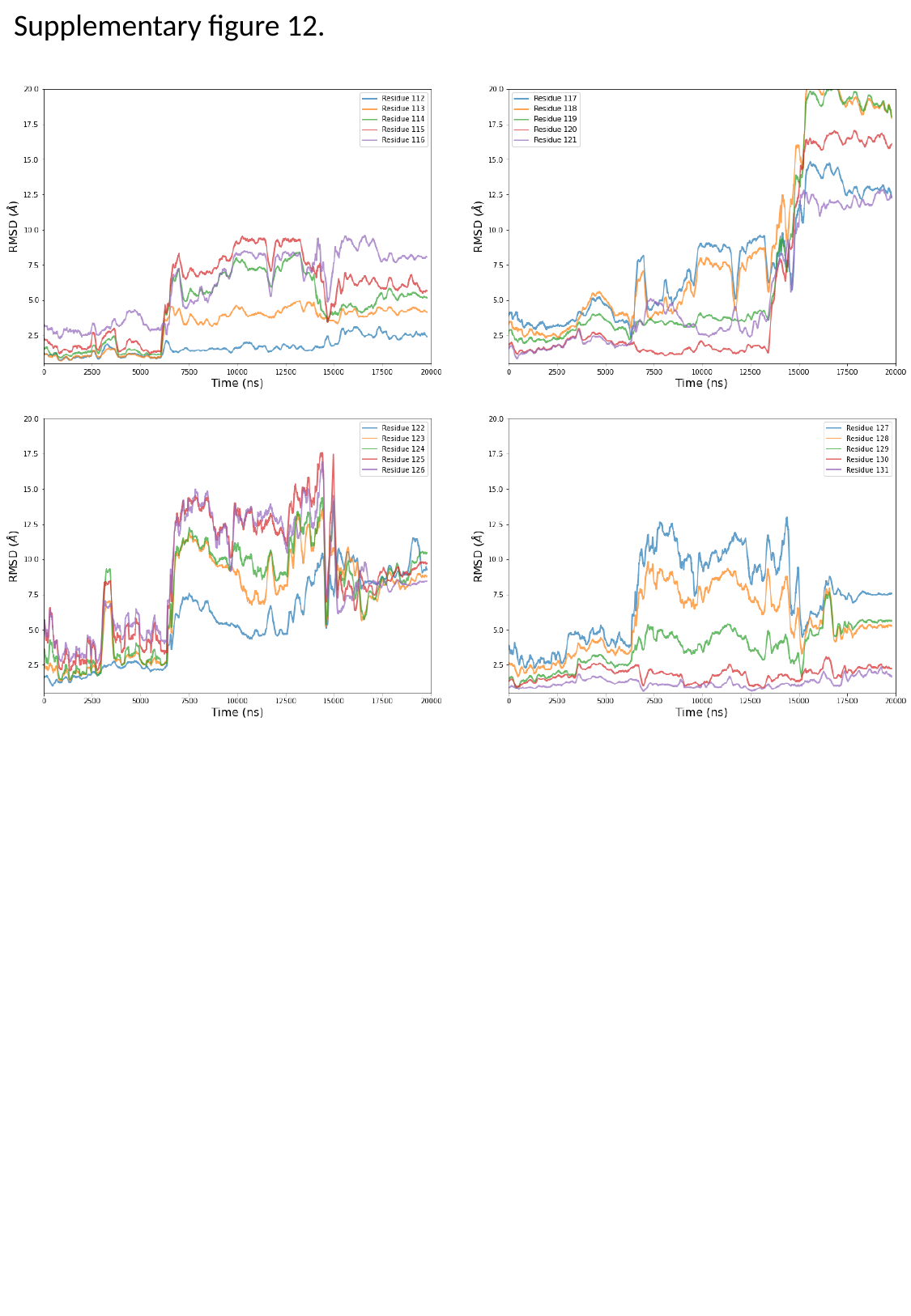

Supplementary figure 12.

### supplementary figure 13

#### Slide 1
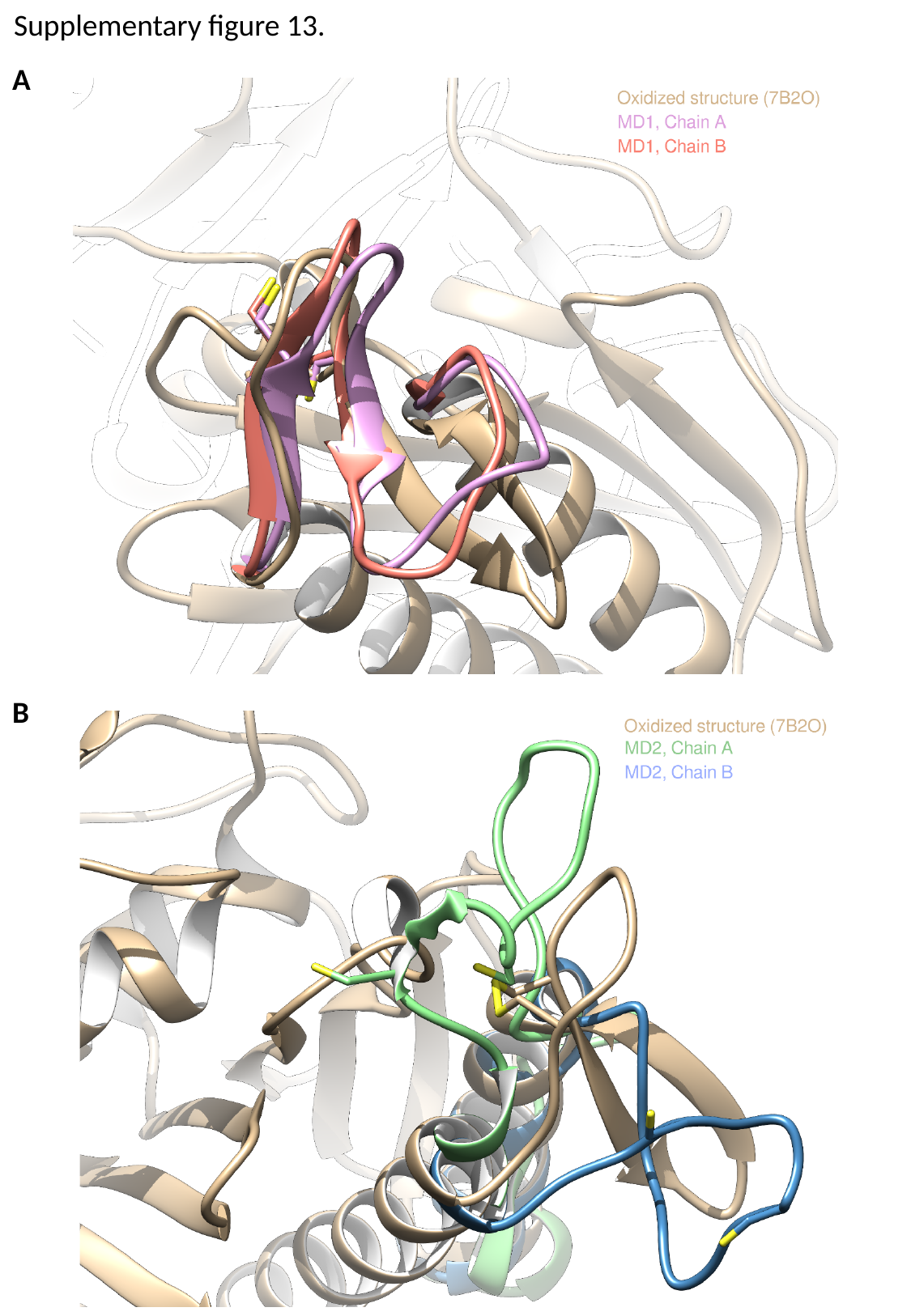

Supplementary figure 13.
A
B

### supplementary figure 14

#### Slide 1
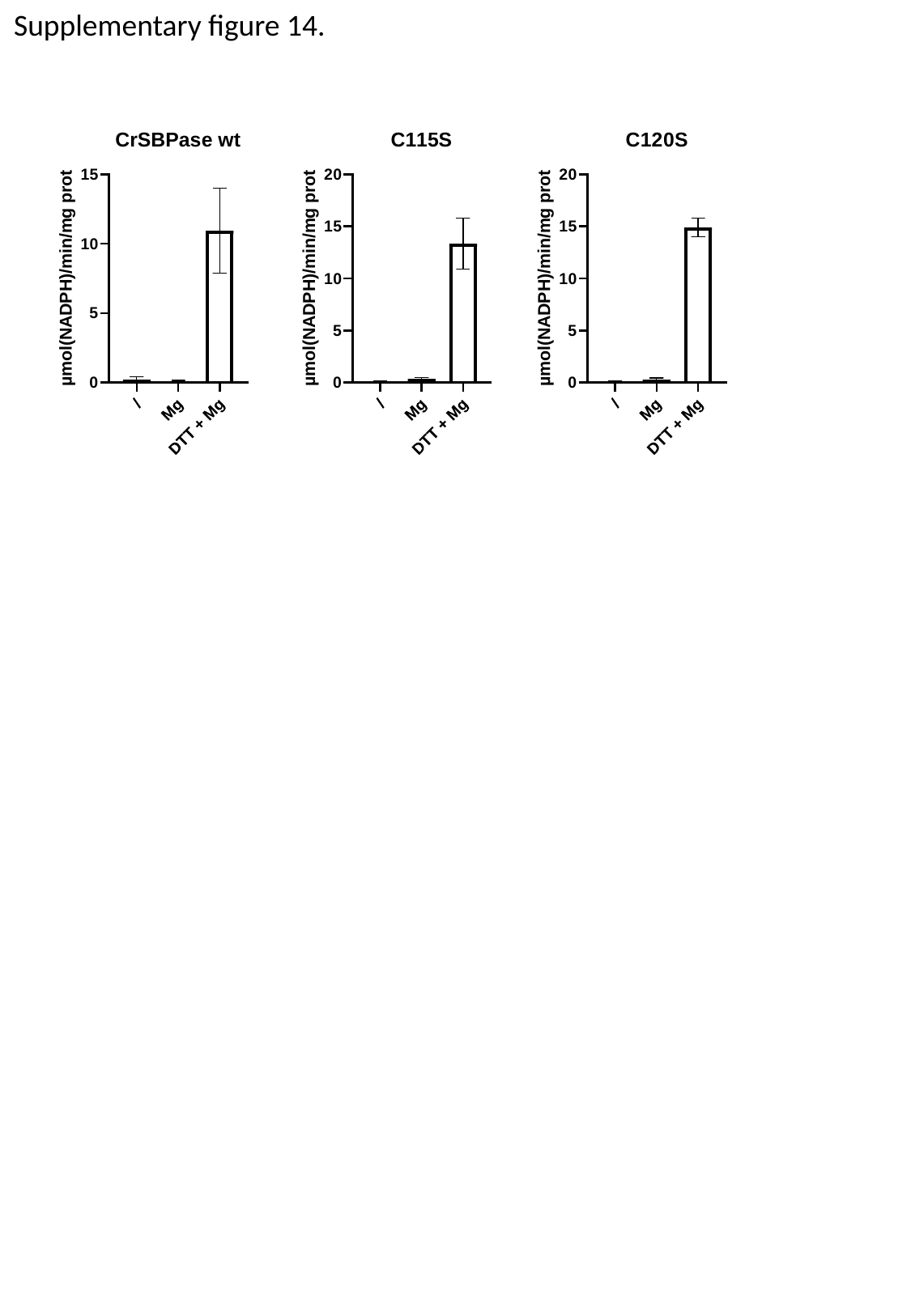

Supplementary figure 14.
