## supplementary figure 5 for "Structure of the Calvin-Benson-Bassham sedoheptulose-1,7-bisphosphatase from the model microalga *Chlamydomonas reinhardtii*"

### Slide 1
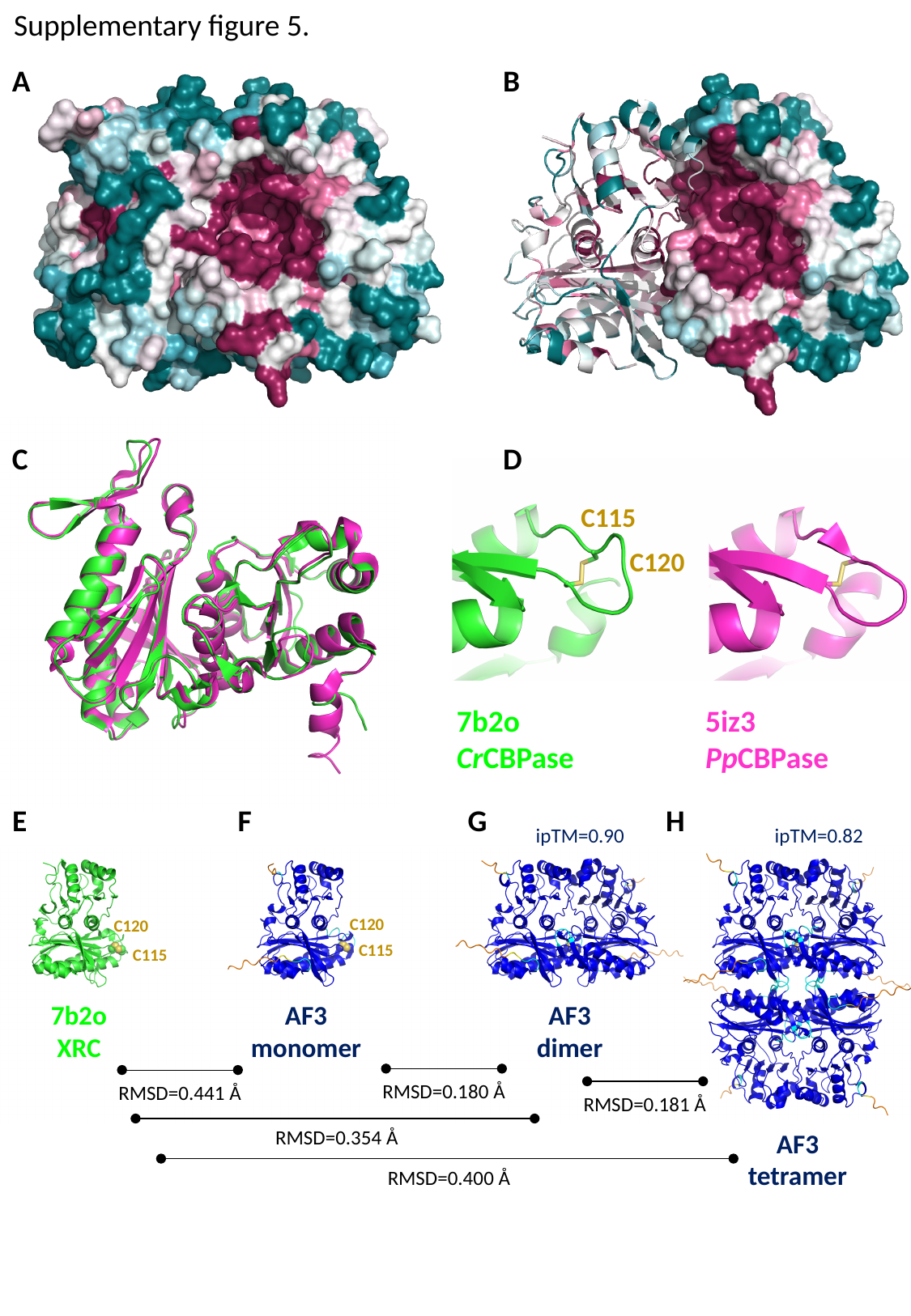

Supplementary figure 5.
A
B
C
D
C115
C120
5iz3
PpCBPase
7b2o
CrCBPase
H
E
F
G
ipTM=0.90
ipTM=0.82
C120
C120
C115
C115
AF3
dimer
7b2o
XRC
AF3
monomer
RMSD=0.180 Å
RMSD=0.441 Å
RMSD=0.181 Å
RMSD=0.354 Å
AF3
tetramer
RMSD=0.400 Å
