## supplementary links for "Structure of the Calvin-Benson-Bassham sedoheptulose-1,7-bisphosphatase from the model microalga *Chlamydomonas reinhardtii*"

**Movies of MD2 in the reduced state for Chain A and Chain B.**

In grey is shown the crystallized structure in the oxidized state. Each second of the video represents 94ns of the MD simulation.

Chain A (434.4 Mb)

<https://dropsu.sorbonne-universite.fr/s/5JDRT6med35y2Jf>

Chain B (441.2 Mb)

<https://dropsu.sorbonne-universite.fr/s/ttDdRqye5ADe9GA>
